# The synthetically produced predator odor 2,5-dihydro-2,4,5-trimethylthiazoline increases alcohol self-administration and alters basolateral amygdala response to alcohol in rats

**DOI:** 10.1101/2020.01.10.901736

**Authors:** Viren H. Makhijani, Janay P. Franklin, Kalynn Van Voorhies, Brayden Fortino, Joyce Besheer

## Abstract

Post-traumatic stress disorder (PTSD) is a psychiatric illness that can increase the risk for developing an alcohol use disorder (AUD). While clinical data has been useful in identifying similarities in the neurobiological bases of these disorders, preclinical models are essential for understanding the mechanism(s) by which PTSD increases the risk of developing AUD. The purpose of these studies was to examine if exposure of male Long-Evans rats to the synthetically produced predator odor 2,5-dihydro-2,4,5-trimethylthiazoline (TMT) would increase alcohol self-administration, potentially by facilitating transfer of salience towards cues, and alter neuronal response to alcohol as measured by the immediate early gene c-Fos. In Experiment 1 rats exposed to repeated (4x) TMT showed reductions in goal-tracking behavior in Pavlovian conditioned approach, and increases in alcohol self-administration. In Experiment 2 rats exposed to repeated TMT showed blunted basolateral amygdala c-Fos response to alcohol, and increased correlation between medial prefrontal cortex and amygdala subregions. In Experiment 3 rats exposed to single, but not repeated TMT showed increases in alcohol self-administration, and no change in anxiety-like behavior or hyperarousal. In Experiment 4, rats showed no habituation of corticosterone response after 4 TMT exposures. In summary, exposure of male rats to TMT can cause escalations in alcohol self-administration, reductions in goal-tracking behavior, and reduction in BLA response to alcohol. These studies outline and utilize a novel preclinical model that can be used to further neurobiological understanding of the relationship between PTSD and AUD.

## 1. Introduction

Alcohol is the one of the most widely consumed and abused psychoactive substances in the United States, with approximately 56% of adults having drank alcohol in the past month and 6% of adults having an alcohol use disorder (SAMHSA, 2018).One risk factor for development of alcohol use disorder (AUD) is comorbid post-traumatic stress disorder (PTSD), which epidemiological studies have estimated nearly triples the risk of developing AUD (Kessler et al., 1997, Jacobsen et al., 2001, Shorter et al., 2015).

PTSD stems from a person witnessing or experiencing a traumatic or life-threatening event. While the symptoms of PTSD have been well-documented (intrusion, avoidance, negative alterations in cognition and mood, alterations in arousal), our understanding of the neurobiological underpinnings are still emerging. Neuroimaging studies have characterized dysregulation of multiple brain regions in PTSD, including hyper-responsivity of the amygdala to stressful stimuli and blunted medial prefrontal cortex (mPFC) top-down control of attentional and amygdala function (Liberzon and Sripada, 2008, Fenster et al., 2018). Unsurprisingly, dysregulation in these brain circuits are also heavily implicated in AUD (Koob and Volkow, 2016, Blaine and Sinha, 2017), and this overlap is thought, in part, to underlie comorbidity between PTSD and AUD (Gilpin and Weiner, 2017).

Animal models play an important role in experimentally probing the relationship between ‘trauma’ exposure and increased alcohol drinking, and a number of animal models of PTSD have been shown to increase alcohol drinking. For example, stress-enhanced fear learning (SEFL) is a model that involves exposing a rat to 15 shocks in one context, which enhances later fear learning in another context (Rau and Fanselow, 2009). Rats that undergo SEFL show increased alcohol consumption in a two-bottle choice protocol for over 120 days (Meyer et al., 2013). Another model of PTSD is exposure to a predator or predator odor. Exposure to bobcat urine (Edwards et al., 2013), soiled cat litter (Manjoch et al., 2016), or a live un-neutered cat in conjunction with social instability (Zoladz et al., 2018) have been shown to increase alcohol drinking in rats for 20, 21, and 7 days respectively in self-administration or two-bottle choice paradigms. Similarly, exposure to dirty rat bedding sex-dependently increases homecage alcohol consumption in mice with a history of binge alcohol consumption (Finn et al., 2018). These studies have begun to advance our knowledge of the link between PTSD and excessive alcohol drinking by contributing models that can be used to identify causal neuromolecular and brain circuitry changes.

The first goal of the present work was to determine if repeated exposure of male rats to the synthetically produced predator odor (PO) 2,5-dihydro-2,4,5-trimethylthiazoline (TMT; an extract of fox feces) would increase reward cue salience and alcohol self-administration. As maladaptive response to cues is a hallmark of both PTSD (APA, 2013, VanElzakker et al., 2014) and AUD (Seo and Sinha, 2014, Valyear et al., 2017) we hypothesized that increases in cue salience may be related to increases in alcohol self-administration. Next, this study quantified alterations in mPFC and amygdala neuronal response to alcohol following TMT-exposure using c-Fos immunoreactivity. Last, this study examined if a single TMT exposure was sufficient to induce changes in alcohol self-administration, and if acute physiological and endocrine response to TMT habituated across repeated exposures.

## 2. Materials and Methods

### 2.1. Animals

96 male Long-Evans rats (Envigo-Harlan, Indianapolis, IN) arrived at 7 weeks old and were single housed under a 12 hour light/dark cycle (7:00 am/pm). Prior to all experiments rats were handled for 1-2 minutes across 7 days. All experiments were conducted during the light cycle, with the exception of TMT exposures which occurred at the start of the dark cycle (7:00 - 7:30 pm). Animals were under the care of the veterinary staff of the UNC-Chapel Hill Division of Comparative Medicine. All procedures were carried out in accordance with the NIH Guide for Care and Use of Laboratory Animals, and institutional guidelines. All protocols were approved by the UNC Institutional Animal Care and Use Committee (IACUC). UNC-Chapel Hill is accredited by the Association for Assessment and Accreditation of Laboratory Animal Care (AAALAC).

### 2.2. Experiment 1: Effect of TMT exposure on Pavlovian conditioned approach (Figure 1A)

The goal of this experiment was to assess if exposure to TMT would later increase the transfer of incentive salience towards reward cues using a Pavlovian conditioned approach (PCA) task. PCA was assessed in conditioning chambers (Med Associates, St. Alban, VT) located within sound-attenuating cabinets equipped with an exhaust fan to provide ventilation and mask outside noise. Chambers were equipped with a retractable lever on the back left side of the chamber and a cue light was located above the lever. Next to the lever was a port containing an infrared beam to detect entries and a liquid receptacle connected to a syringe pump for delivery of sucrose (20%, w/v).

#### 2.2.1. PCA training

One day prior to PCA, rats underwent a pretraining session to familiarize them with the sucrose reinforcer (Fig 1A). The pretraining session consisted of 25 non-contingent presentations of 20% sucrose (0.1 mL delivered by a syringe pump across 1.66 s) into the liquid receptacle on a 90 second variable interval (30 – 150 s) schedule. Across the next 8 days rats were trained on PCA. PCA sessions consisted of 25 trials on a 90 second variable interval schedule. Trials consisted of a 10 second presentation of a lever and cue light (conditioned stimulus, CS) during which lever presses and port entries were measured. After 10 seconds the lever retracted, the cue light was turned off, and 0.1 mL of 20% sucrose (unconditioned stimulus, US) was delivered into the liquid receptacle port.

**Fig. 1.**
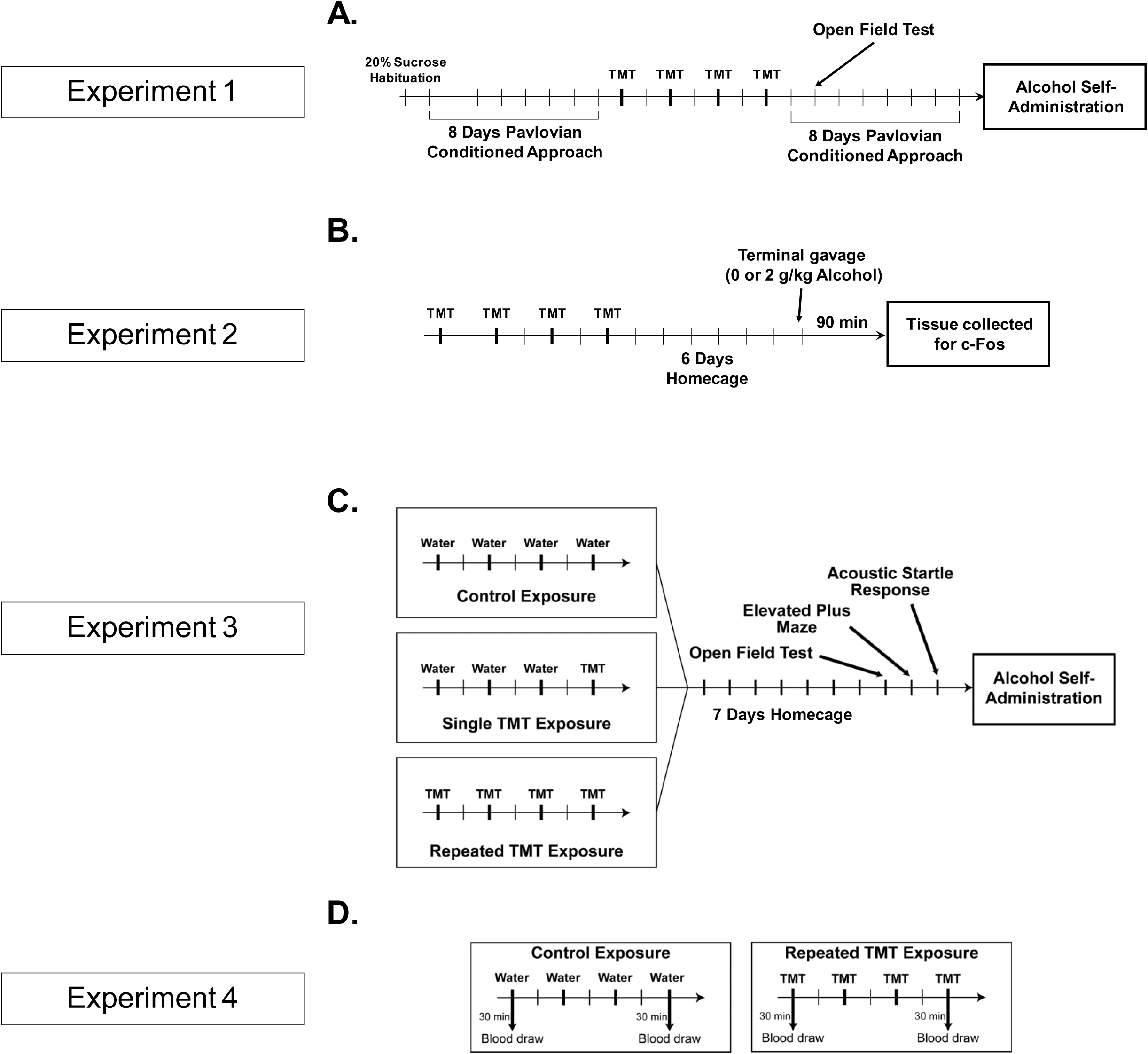
Timelines for Experiments 1-4. (A) In Experiment 1, male rats (n = 6/group) were trained on PCA for 8 sessions prior to 4 TMT exposures. Rats then returned to PCA for 8 additional sessions and underwent an open field test 48 hours after the final TMT exposure. After PCA rats were trained to self-administer alcohol. (B) In Experiment 2, male rats (n = 6/dose control group and 10/dose TMT group) were exposed to TMT 4 times. 7 days later, rats received 2 g/kg alcohol or water (i.g.) and were sacrificed 90 minutes later to examine brain regional expression of c-Fos. (C) In Experiment 3, male rats (n = 12/group) were exposed 0, 1, or 4 times to TMT and examined for anxiety-like behavior and hyperarousal 7 days later. Rats were then trained to self-administer alcohol. (D). In Experiment 4, male rats (n = 8/group) were exposed to TMT 4 times and blood was drawn 30 minutes post-exposure on days 1 and 4 for analysis of plasma corticosterone.

#### 2.2.2. TMT exposures

After 8 PCA sessions rats (n = 6/group) underwent TMT exposures. Animals were transported from the vivarium to a well-ventilated room at the onset of the dark cycle (7:00pm) and placed into clean Plexiglas chambers (30.5 cm × 18 cm × 18 cm) with a piece of filter paper affixed to the lid. 10 µL of either water (Control group) or TMT was pipetted onto the filter paper prior to closing the chamber lids. (Note: Control animals were always exposed before TMT animals to avoid any contact with the TMT odor). After 10 minutes of exposure, rats were returned to the vivarium. This TMT exposure process was repeated 4 times across one week (i.e., every other day; on the intervening days rats remained in the home cage undisturbed). PCA training was withheld during this TMT exposure period.

Following the final TMT exposure, rats underwent 8 additional PCA sessions. Before the second PCA session rats underwent an open field test (between 08:00 am - 12:00 pm) to assess anxiety-like behavior and locomotor behavior. Rats were transported in the home cage and were allowed to habituate to the testing room for 20 minutes. Rats were placed in the center of the open field arena (43 cm × 43 cm; Med Associates, St. Albans, VT) and locomotor activity was recorded by 32 orthogonal infrared beams using Activity Monitor software (Med Associates) for 10 min.

#### 2.2.3. Alcohol self-administration

After completion of the PCA phase of the experiment, rats began alcohol self-administration. The alcohol self-administration chambers were the same chambers used for PCA. However, the configuration of the chamber was such that there was an additional lever and cue light on the opposite side of the chamber (i.e. two levers and two cue lights total, one set on the left side of the chamber, by the fluid port, and one on the right side). Additionally, general locomotor activity during the self-administration session was measured with 4 parallel infrared beams across the chamber floor. Total beam breaks across the session were collected and this number was divided by the session length (30 min) to determine locomotor rate (beam breaks/min). Rats were trained to self-administer a 15% (v/v) alcohol + 2% (w/v) sucrose solution (15A/2S) on a fixed ratio 2 (FR2) schedule of reinforcement in 30 minute sessions, five days a week (M-F) via sucrose fading as described in Makhijani et al., 2018. Sucrose fading began with self-administration of 10% sucrose (10S; 2-3 sessions), then 2A/10S (1-2 sessions), 5A/10S (1 session), 10A/10S (2 sessions), 10A/5S (1 session), 15A/5S (2 sessions), and 15A/2S (2 sessions). Finally there were 5 sessions of 15A reinforcer after which the reinforcer returned to 15A/2S for the duration of training (maintenance phase). A sweetened alcohol reinforcer was used as we find this results in stable alcohol self-administration in these long-term studies (Randall et al., 2017, Jaramillo et al., 2018, Makhijani et al., 2018).

### 2.3. Experiment 2: Effect of TMT exposure on neuronal response to alcohol (Figure 1B)

The goal of this experiment was to examine potential changes in neuronal response to acute alcohol following predator odor stress. Rats (n = 6/dose control group and 10/dose TMT group) experienced the repeated TMT exposure protocol as in Experiment 1. Rats remained undisturbed in the home cage for the next 6 days. On the final test day (7 days after the final TMT exposure), rats received an alcohol injection (2 g/kg alcohol or equivalent volume of water; intragastrically, IG). 90 minutes later, rats were sacrificed by pentobarbital anesthesia (100 mg/kg) prior to perfusion with 0.1M PBS (4°C, pH = 7.4) followed by 4% paraformaldehyde (PFA; 4°C, pH = 7.4). Brains were extracted and stored in 4% PFA for 24h at 4°C before being rinsed with 0.1M PBS and transferred to 30% sucrose in 0.1M PBS for at least 7 days. Brains were sliced on a freezing microtome into 40 µm coronal sections which were stored at −20°C in cryoprotectant until immunohistochemistry (IHC).

#### 2.3.1. c-Fos IHC and quantification of immunoreactivity (IR)

Free-floating coronal sections were rinsed in 0.1M PBS before quenching of endogenous peroxidases with a 5 minute wash in 1% H_2_O_2_. Sections were then blocked with 3% normal goat serum (NGS; Vector Labs, Burlingame, CA) in 0.3% Triton X-100 for 2 hours before being incubated in rabbit anti-c-Fos antibody (1:4000 in 3% NGS + 0.1% Triton; Synaptic Systems, Goettingen, Germany; Lot# 226003/3-45) for 16 hours at 4°C. Sections were incubated with biotinylated goat anti-rabbit secondary antibody (1:200 in 3% NGS + 0.1% Triton X-100; Vector Labs, Burlingame, CA) followed by Vectastain Elite ABC HRP (Vector labs, Burlingame, CA). Finally, sections were treated with diaminobenzidine (Sigma-Aldrich, St. Louis, MO) and mounted on slides for imaging.

Images were taken with an Olympus CX41 light microscope (Olympus America, Center Valley, PA) and analyzed utilizing Image-Pro Premier image analysis software (Media Cybernetics, Rockville, MD). Immunoreactivity data (c-Fos-positive pixels/mm^2^ ) were acquired from a minimum of 2 sections/brain region/animal, by an experimenter blind to group assignment, and the data were averaged to obtain a single value per subject. The regions examined were the prelimbic cortex (PL; AP +3.7 to +3.0), infralimbic cortex (IL; AP +3.7 to +3.0), central amygdala (CeA; AP −1.9 to −2.8), and basolateral amygdala (BLA; AP −1.9 to −2.8).

### 2.4. Experiment 3: Comparison of single and repeated TMT exposure on behavior and alcohol self-administration (Figure 1C)

The goal of this experiment was to determine the consequences of a single vs. repeated TMT exposure on alcohol self-administration. Rats were assigned to one of the following groups (n = 12/group): Control group – rats underwent 4 water exposures; Single TMT (sTMT) group – rats underwent 3 water exposures and a single TMT exposure on the final session; Repeated TMT (rTMT) group – rats underwent 4 TMT exposures. These exposures were every other day for 1 week as described in Experiment 1 (2.2.2.). In addition, the number of fecal boli in the test chambers following TMT exposure was counted and presented as part of Experiment 4. Seven days later, rats were tested in the open field as described in Experiment 1 (2.2.2.), the elevated plus maze, and the acoustic startle test. Rats underwent each test in that order on consecutive days. For the elevated plus maze, rats were placed into the closed arm of the maze (arm dimensions 50 cm × 10 cm; maze elevation 50 cm; Stoelting, Wood Dale, IL) for a 5 minute test under red lighting conditions. Movement was tracked using a ceiling mounted camera (Logitech C615; Logitech, Lausanne, Switzerland) and analyzed by AnyMaze software (Stoelting). For the acoustic startle test, rats were placed in an SR-LAB animal enclosure (San Diego Instruments, San Diego, CA) and testing consisted of a 5 minute habituation to background white noise (65dB, matched to ambient noise), followed by 30 startle trials (40 msec presentation of 110 dB white noise) with a 30 second inter-trial interval. Startle amplitude was measured with a high-accuracy accelerometer mounted under the animal enclosure and analyzed with SR-LAB software (San Diego Instruments). Sessions were approximately 20 min in total. The day following the acoustic startle test, rats began alcohol self-administration as described in Experiment 1 (2.2.3.).

### 2.5. Experiment 4: Measuring plasma corticosterone response to repeated TMT exposure (Figure 1D)

In order to measure the effect of TMT exposure on plasma corticosterone levels, rats underwent either 4 water exposures or 4 TMT exposures across one week as described in section 2.2.2. Tail blood was collected 30 minutes after exposure on days 1 and 4 for analysis of plasma corticosterone. Blood was collected into heparinized tubes and immediately centrifuged at 4°C for 5 minutes at 2000 rcf. Plasma supernatant was then collected and stored at −80°C until analysis. 5 µL plasma samples were then analyzed in duplicate using a commercially available colorimetric EIA kit (ArborAssays, Ann Arbor, MI) according to the manufacturer’s instructions.

### 2.6. Reagents

97% purity 2,5-dihydro-2,4,5-trimethylthiazoline (TMT) was purchased from SRQ Bio (Sarasota, FL). Alcohol (95% (v/v); Pharmaco-AAPER, Shelbyville, KY) and sucrose (Great Value, Bentonville, AR) were diluted with tap water for all PCA and self-administration sessions.

### 2.7. Data analysis

#### 2.7.1. PCA

PCA behavior is represented by cumulative number of lever presses across all trials, cumulative port entry elevation score across all trials (port entries during 10s CS – port entries in 10s preceding CS), and latency to port entry and lever press (from CS onset). PCA behavior is also summarized by a PCA index, which takes into account responses, latency to response, and probability of response during trial as in (Fitzpatrick et al., 2019). PCA index was calculated by an evenly weighted average of response bias ((lever presses – port entry elevation score)/(lever presses + port entry elevation score)), latency score ((latency to port entry– latency to lever press)/10s), and probability difference (probability of lever press in trial – probability of port entry). Animals with a PCA index below −0.5 are considered “Goal-Trackers”, an index above 0.5 indicates a “Sign-Tracker”, and animals in between are considered “Intermediate Responders” (Fitzpatrick and Morrow, 2016, Fitzpatrick et al., 2019). PCA behavior after TMT exposure was analyzed by two-way repeated measures analysis of variance (RM-ANOVA) with TMT exposure as the between-subjects factor and session as the within-subjects factor.

#### 2.7.2. Self-administration

For the sucrose fading phase of alcohol self-administration, alcohol lever responses, inactive lever responses, and locomotor rate are represented as averages across each reinforcer. These data were examined by two-way RM-ANOVA with TMT exposure as a between-subjects factor and reinforcer as a within-subjects factor. Total alcohol intake (g/kg) during the sucrose fading and maintenance phases of self-administration is shown as cumulative alcohol consumed by each animal across the study (approximated based on body weight and number of reinforcers delivered for each session) and compared by t-test for Experiment 1, and one-way ANOVA for Experiment 3. For maintenance sessions of alcohol self-administration: alcohol lever responses, inactive lever responses, and locomotion are presented as 3-session averages. These data were analyzed by two-way RM-ANOVA with TMT exposure as a between-subjects factor, and session as a within-subjects factor.

#### 2.7.3. Other measures

c-Fos immunoreactivity was analyzed by two-way ANOVA with TMT exposure and alcohol treatment as between-subjects factors. Correlations between brain regional c-Fos expression were evaluated by Pearson’s correlation coefficient. Measures from the open field test in Experiment 1 were analyzed by t-test. Correlations between post-TMT change in PCA index and total alcohol consumption were evaluated by Pearson’s correlation coefficient. Measures from the behavioral screens (% center time in open field [center defined as middle 20 cm × 20 cm of open field], total distance traveled in open field, % open arm time in elevated plus maze, total distance traveled in elevated plus maze, peak startle response, startle habituation index) in Experiment 3 were analyzed by one-way ANOVA with TMT exposure as the between-subjects factor. The formula for startle habituation index is: ((Average Peak Startle in First 5 Trials) – (Average Peak Startle in Last 5 Trials))/(Average Peak Startle in First 5 Trials). Plasma corticosterone levels (ng/ml) following TMT were analyzed by two-way RM-ANOVA with TMT exposure as a between-subjects factor and exposure day as a within-subjects factor. Fecal boli was analyzed by two-way RM-ANOVA with TMT exposure as a between-subjects factor, and exposure day as the within-subjects factor. All data is represented as mean ± SEM. For all analyses significance was set at p < 0.05.

## 3. Results

### 3.1. Experiment 1: Effect of TMT exposure on Pavlovian conditioned approach

The purpose of Experiment 1 was to determine if TMT exposure would increase transfer of salience towards reward cues (sign-tracking), and if increased cue salience/sign-tracking behavior could predict increased alcohol self-administration.

#### 3.1.1. Repeated TMT exposure reduces goal-tracking behavior towards a reward cue

Male rats underwent 8 days of PCA training, after which animals were counterbalanced into groups by PCA index (measured on day 8) and assigned to either the repeated TMT or Water group. Following TMT exposure, animals showed no change in lever presses (Fig 2A), but showed significantly lower port entry elevation scores than the control group (Fig 2B, F(1,10) = 5.18, p = 0.046), with no significant main effect of session or TMT by session interaction. There were no main effects of TMT or session on latency to first lever press (Fig 2C) and there was a trend for a main effect of TMT on latency to first port entry (Fig 2D, F(1,10) = 4.05, p = 0.072), with no main effect of session or TMT by session interaction. There were no main effects of TMT, session or interaction on PCA index (Fig 2E). Additionally, while all rats performed sign-tracking behavior (lever presses) indicating transfer of salience to the lever cue, no rats in either group met the classification of sign-tracker (PCA index over 0.5). Both groups had 2 goal-trackers (PCA index under −0.5) and 4 intermediate responders (PCA index between −0.5 and 0.5). These findings indicate that TMT exposure did not alter sign-tracking behavior but did reduce goal-tracking behavior. In the open field test there was no effect of TMT exposure on distance travelled or percent time in the center of the open field (Table 1), indicating the lack of a change in general locomotion and anxiety-like behavior.

**Fig. 2.**
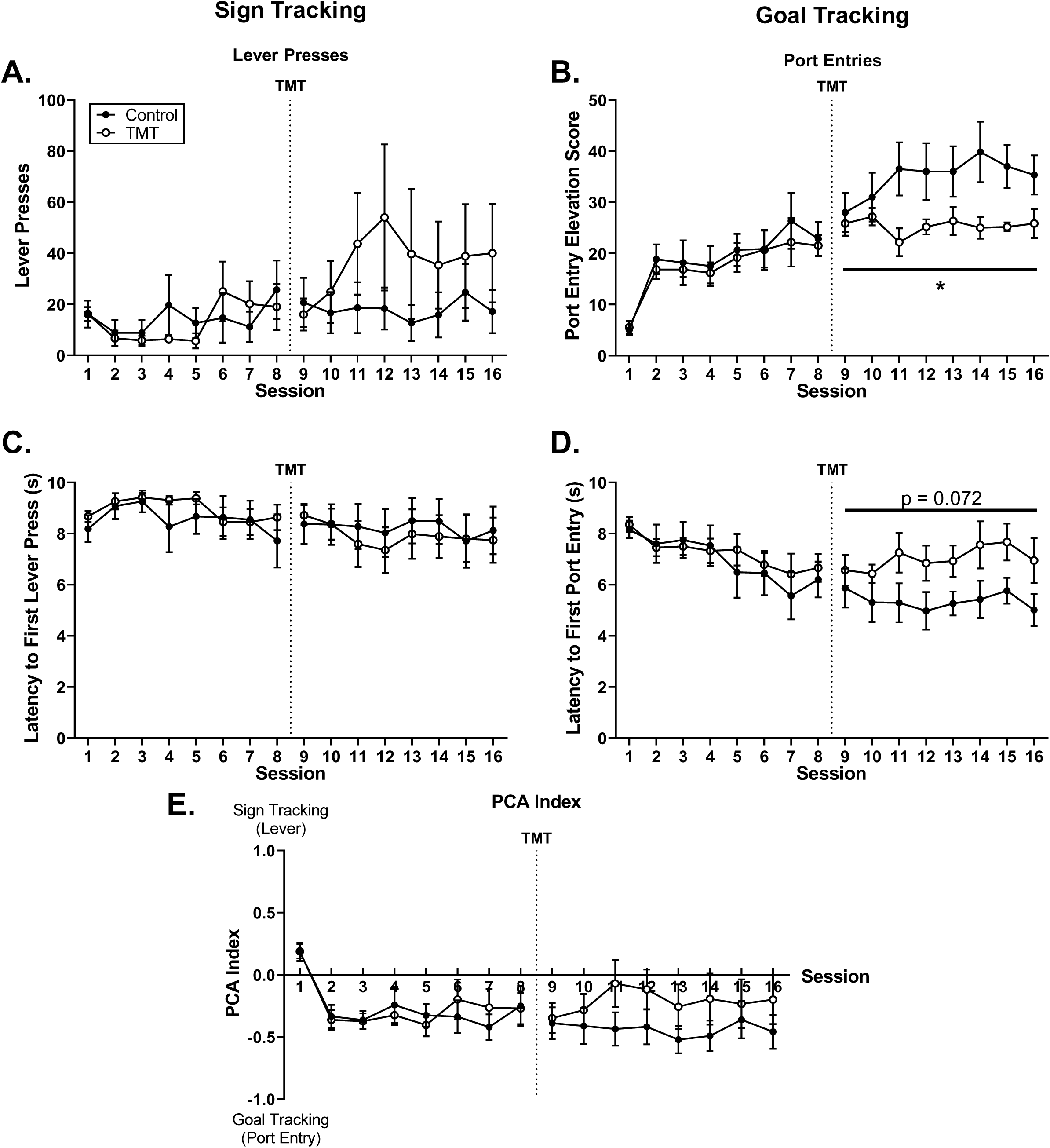
Effects of TMT exposure on PCA behavior. (A&B) Rats exposed to TMT showed no change in lever presses (sign-tracking) and significant reductions in port-entry elevation score (goal-tracking) compared to controls. (C&D) Rats exposed to TMT showed no change in latency to first lever press, and a trend for increased latency to first port-entry as compared to controls. (E) PCA indices for TMT exposed rats did not differ significantly from controls. Dotted line represents 4 TMT exposures across 7 days, between PCA sessions 8 and 9. * - p < 0.05 versus control.

**Table 1.** Behavioral data from Experiments 1 and 3.

| Group | Open Field |  | Elevated Plus Maze |  | Acoustic Startle |  |
| --- | --- | --- | --- | --- | --- | --- |
|  | Center Time (%) | Total Distance (cm) | Open Arm Time (%) | Total Distance (m) | Average Peak Startle (mV) | Habituation Index (%) |
| **Experiment 1** |  |  |  |  |  |  |
| Control | 13.6 ± 2.9 | 1633.7 ± 306.9 | - | - | - | - |
| TMT | 20.4 ± 1.4 | 1283.3 ± 435.3 | - | - | - | - |
| **Experiment 3** |  |  |  |  |  |  |
| Control | 12.8 ± 1.2 | 1547.2 ± 84.0 | 4.3 ± 2.6 | 10.2 ± 1.1 | 650.2 ± 197.2 | 0.3 ± 15.3 |
| sTMT | 12.5 ± 1.0 | 1671.3 ± 109.6 | 5.7 ± 3.6 | 8.7 ± 0.9 | 376.6 ± 91.9 | 9.1 ± 19.2 |
| rTMT | 10.4 ± 1.3 | 1495.7 ± 84.8 | 7.5 ± 2.5 | 9.5 ± 0.7 | 358.2 ± 55.3 | 3.0 ± 20.4 |

#### 3.1.2. Rats with a history of repeated TMT exposure show increased alcohol self-administration

Two-way RM ANOVA across sucrose fading showed main effects of reinforcer and TMT on alcohol lever responses with no interaction (Fig 3A; F _Reinforcer_(7,70) = 36.7, p < 0.001; F_TMT_(1,10) = 6.14, p = 0.033) indicating greater alcohol lever responses in the TMT group than the Control group. The TMT-exposed animals also had significantly higher total alcohol intake across sucrose fading (Fig 3B; t(10) = 2.59, p = 0.027). There were no main effects of TMT or reinforcer on inactive lever responses or locomotor rate (Table 2).

**Fig. 3.**
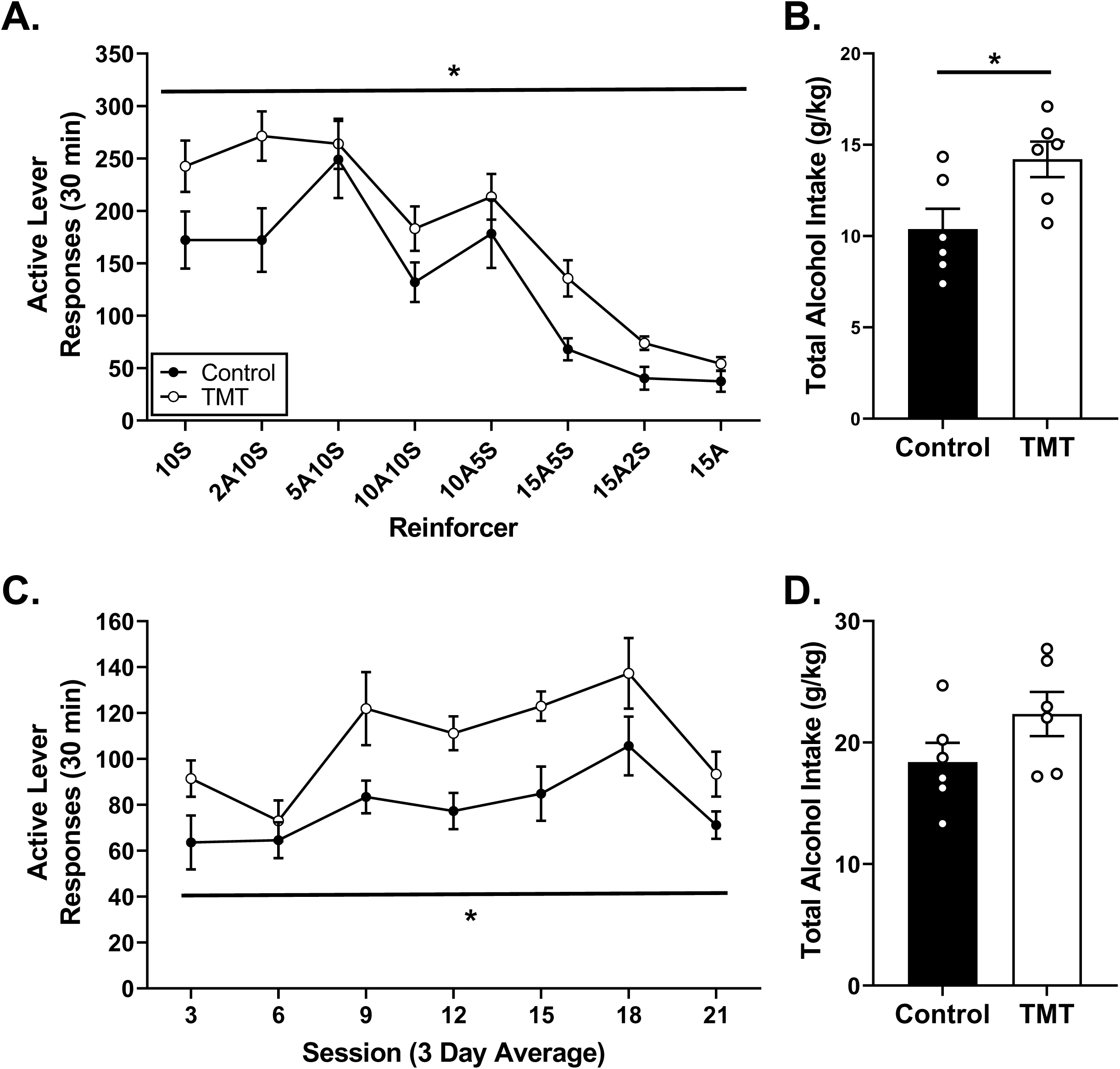
Acquisition and maintenance of alcohol self-administration following TMT exposure in Experiment 1. (A) Rats exposed to TMT showed increased active lever responses across sucrose fading compared to controls. (B) TMT-exposed rats consumed significantly more alcohol than controls across sucrose fading. (C) TMT-exposed rats showed increased active lever responses during maintenance compared to controls. (D) There was no significant difference in total alcohol intake across the maintenance phase between TMT-exposed animals and controls. * - p < 0.05 versus control.

**Table 2.** Inactive lever responses and locomotor rate from sucrose fading phase in Experiments 1 and 3.

|  |  | **Inactive Lever Responses** |  |  |  |  |  |  |  |
| --- | --- | --- | --- | --- | --- | --- | --- | --- | --- |
| Group |  | 10S | 2A10S | 5A10S | 10A10S | 10A5S | 15A5S | 15A2S | 15A |
| **Experiment 1** |  |  |  |  |  |  |  |  |  |
|  | Control | 0.9 ± 0.3 | 0.8 ± 0.5 | 1.0 ± 0.4 | 1.4 ± 0.5 | 1.2 ± 0.5 | 2.3 ± 1.0 | 1.3 ± 0.7 | 2.5 ± 0.6 |
|  | TMT | 3.1 ± 1.0 | 5.2 ± 2.1 | 6.0 ± 2.1 | 4.4 ± 1.9 | 3.8 ± 1.5 | 5.1 ± 3.0 | 4.2 ± 2.0 | 5.4 ± 2.9 |
| **Experiment 3\*** |  |  |  |  |  |  |  |  |  |
|  | Control | 2.1 ± 0.6 | 2.9 ± 0.8 | 3.5 ± 1.3 | 3.6 ± 0.9 | 2.3 ± 0.9 | 3.0 ± 0.6 | 3.0 ± 0.8 | 2.2 ± 0.6 |
|  | sTMT | 1.9 ± 0.5 | 2.2 ± 0.6 | 3.4 ± 0.7 | 3.2 ± 0.8 | 4.4 ± 1.0 | 2.4 ± 0.7 | 3.3 ± 1.0 | 2.4 ± 0.6 |
|  | rTMT | 3.7 ± 0.9 | 2.7 ± 0.6 | 4.8 ± 1.4 | 3.3 ± 0.7 | 4.3 ± 1.1 | 4.3 ± 1.7 | 3.1 ± 0.8 | 2.6 ± 0.5 |
|  |  | **Locomotor Rate (Beam Breaks/min)** |  |  |  |  |  |  |  |
| Group |  | 10S | 2A10S | 5A10S | 10A10S | 10A5S | 15A5S | 15A2S | 15A |
| **Experiment 1** |  |  |  |  |  |  |  |  |  |
|  | Control | 20.7 ± 3.0 | 21.6 ± 1.3 | 22.6 ± 2.3 | 23.6 ± 1.3 | 26.0 ± 1.2 | 24.5 ± 1.4 | 23.1 ± 1.8 | 24.3 ± 1.4 |
|  | TMT | 28.4 ± 2.0 | 28.0 ± 1.6 | 26.5 ± 2.7 | 27.3 ± 2.1 | 28.7 ± 2.6 | 27.1 ± 2.2 | 25.1 ± 1.9 | 25.5 ± 1.7 |
| **Experiment 3\*** |  |  |  |  |  |  |  |  |  |
|  | Control | 26.7 ± 1.5 | 26.3 ± 1.2 | 27.2 ± 1.6 | 24.7 ± 1.5 | 23.7 ± 1.5 | 23.9 ± 1.1 | 26.5 ± 1.3 | 25.3 ± 1.3 |
|  | sTMT | 29.8 ± 1.3 | 29.0 ± 1.7 | 30.3 ± 2.1 | 26.9 ± 1.1 | 26.8 ± 1.3 | 26.0 ± 0.9 | 28.3 ± 1.5 | 27.0 ± 1.4 |
|  | rTMT | 29.1 ± 1.2 | 29.0 ± 1.4 | 29.9 ± 1.2 | 28.6 ± 0.7 | 29.0 ± 0.8 | 26.4 ± 1.3 | 29.3 ± 1.6 | 27.3 ± 1.4 |

\* - significant main effect of reinforcer $p < 0.05$

During maintenance of alcohol self-administration the TMT group had higher alcohol lever responses than the control group, as indicted by a significant main effect of group (Fig 3C; F(1,10) = 8.11, p = 0.017). There was also a significant main effect of session (F(6,60) = 10.2, p < 0.001) and no interaction. There was no significant effect of TMT exposure on total alcohol intake during the maintenance phase (Fig 3D). There were no main effects of group, session, or interaction for inactive lever responses (Table 3). There was a main effect of session on locomotor rate with reduced locomotion in later sessions (Table 3; F(6,60) = 5.17, p < 0.001), but no main effect of group or session by group interaction.

**Table 3.**
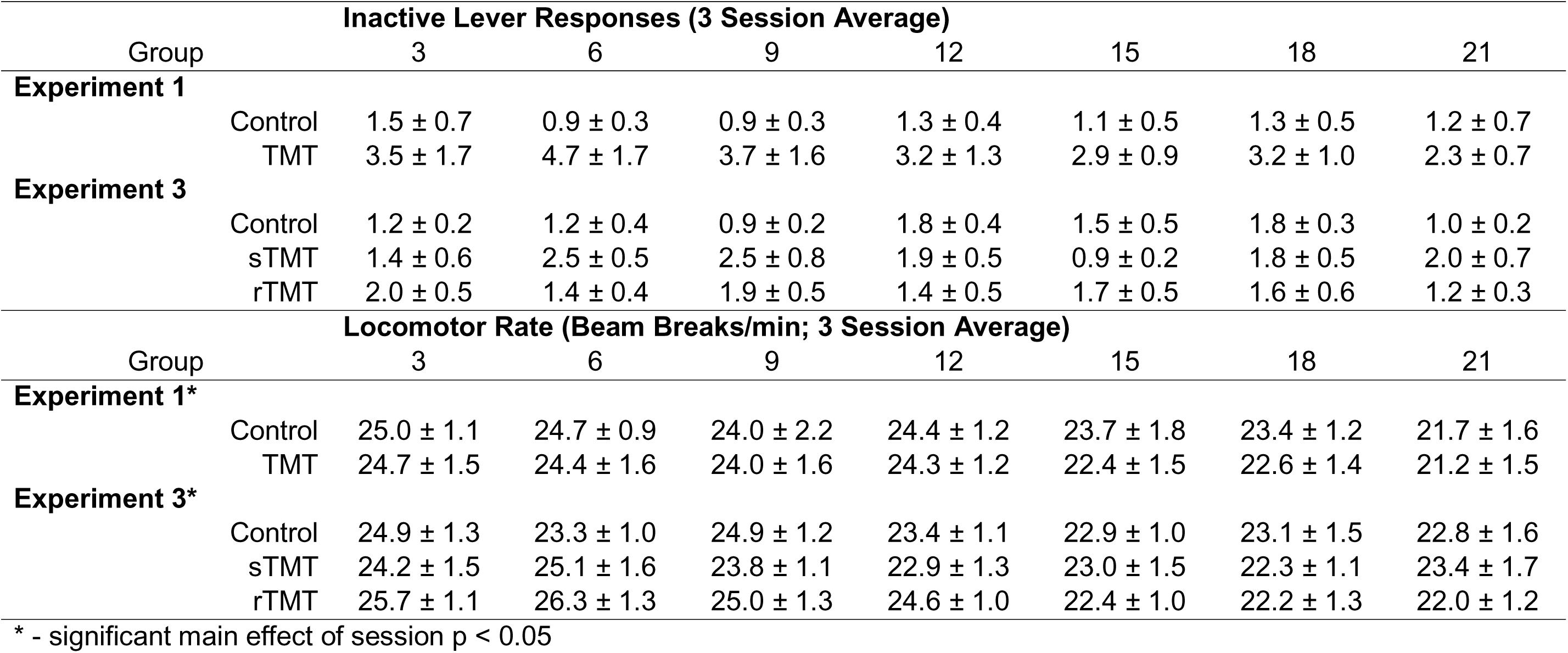
Inactive lever responses and locomotor rate from maintenance phase of self-administration in Experiments 1 and 3.

Together these results show that TMT exposure does not increase transfer of salience to a reward cue, but can reduce goal-tracking behavior. As it was hypothesized transfer of salience to appetitive cues may predict stress induced increases in alcohol consumption, correlation between change in PCA index following TMT exposure (difference between sessions 16 and 8) and total alcohol intake during maintenance self-administration was examined. There were no significant correlations between change in PCA score following TMT exposure and alcohol self-administration in either the control or TMT group (R^2^ = 0.257, R^2^ = 0.001).

### 3.2. Experiment 2: Effect of TMT exposure on neuronal response to alcohol

As rats exposed to repeated TMT showed increased alcohol self-administration, the goal of this experiment was to determine whether neuronal response to alcohol was altered in rats exposed to TMT. Due to experimenter error and tissue damage during sectioning, the animals in the Control + Alcohol group is n = 5 for mPFC regions and n = 4 for amygdala regions.

#### 3.2.1. Repeated TMT exposure blocks alcohol-induced reduction in BLA c-Fos

Following alcohol (2 g/kg), a significant reduction in c-Fos IR was observed in the prelimbic (PL; Fig 4A; F(1,27) = 4.82, p = 0.0367) and infralimbic (IL; Fig 4B; F(1,27) = 5.28, p = 0.030) subregions of the mPFC, with an increase in the central amygdala (CeA; Fig 4C; F(1,25) = 6.29, p = 0.019). There were no main effects of TMT exposure or TMT exposure by alcohol dose interactions.

**Fig. 4.**
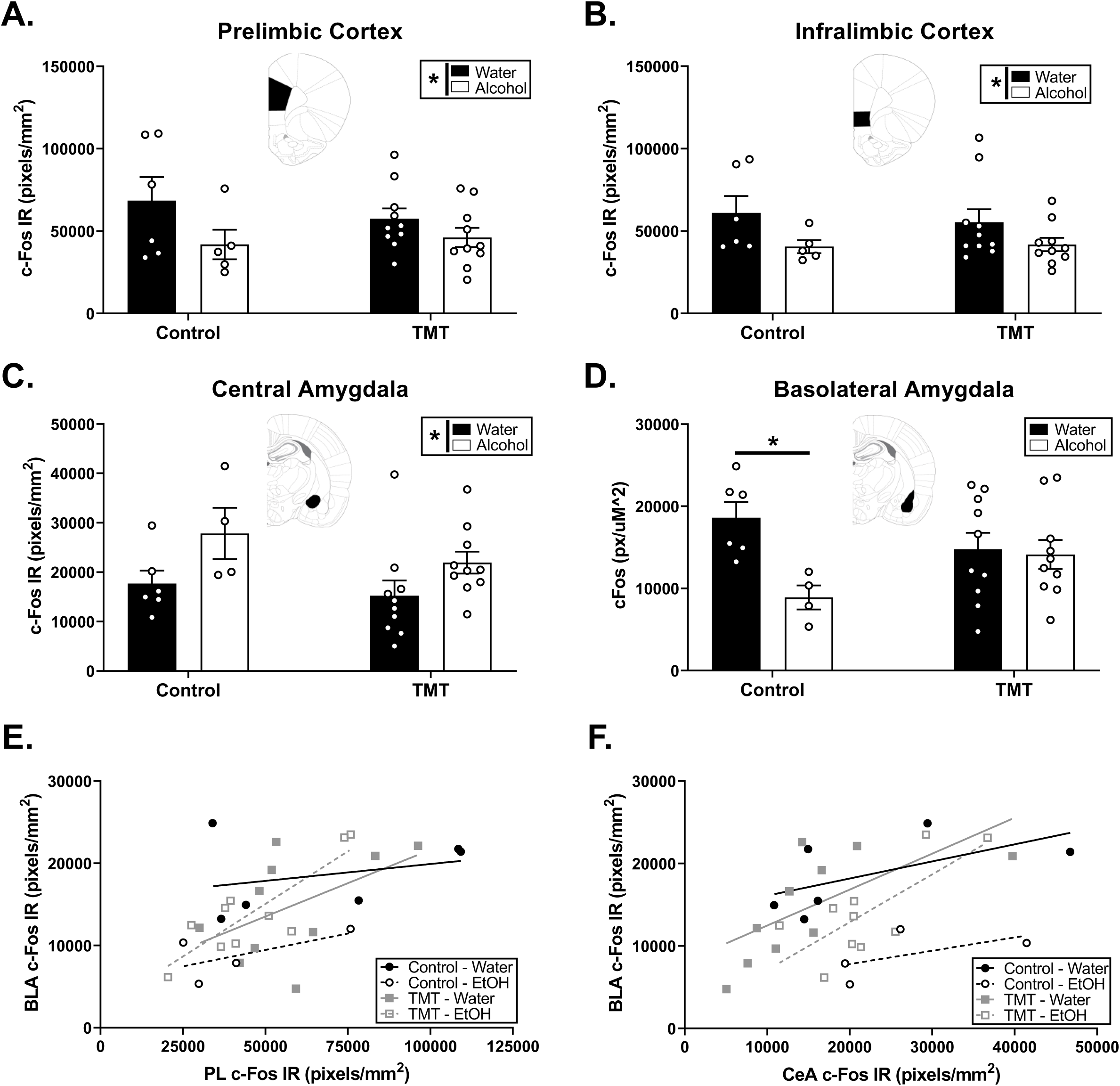
Alcohol induced c-Fos in naïve and TMT-exposed rats. (A, B) Following an alcohol (2 g/kg, IG) injection, there were significant reductions in PL and IL c-Fos IR. (C) Alcohol treated rats showed increased CeA c-Fos IR regardless of TMT history. (D) Alcohol reduces BLA c-Fos IR in naïve, but not TMT-exposed rats. The shaded area on the brain atlas illustrations show the quantified region. * - p < 0.05 versus water.

In the basolateral amygdala (BLA) there was a main effect of alcohol (Fig 4D; F(1,26) = 5.78, p = 0.024) with a significant alcohol by TMT exposure interaction (F(1,26) = 4.48, p = 0.044). Post-hoc analysis showed alcohol treatment significantly reduced BLA c-Fos IR in the control group (p < 0.05), but this effect was blocked in the TMT-exposed group.

Interestingly, the TMT-exposed animals also showed several significant correlations between BLA, CeA, PL, and IL c-Fos IR (Table 4). Notably, the TMT-exposed animals treated with alcohol showed significant correlations between PL and BLA c-Fos IR (Fig 4E; R^2^ = 0.724, p = 0.002) while the Control and TMT + Water groups did not. Both TMT treated groups also showed significant correlations between CeA and BLA c-Fos IR (Fig. 4F; TMT + Water: R^2^ = 0.448, p = 0.034; TMT + EtOH: R^2^ = 0.538, p = 0.016), while neither control group did.

**Table 4.** Correlations between Experiment 2 brain regional c-Fos IR.

| Group | PL – IL | PL – BLA | PL – CeA | IL – BLA | IL – CeA | BLA - CeA |
| --- | --- | --- | --- | --- | --- | --- |
| Control – Water | 0.517 | 0.095 | 0.149 | 0.086 | 0.699* | 0.367 |
| Control – Alcohol | 0.848* | 0.391 | 0.066 (-) | 0.666 | 0.000 | 0.321 |
| TMT – Water | 0.810* | 0.248 | 0.479* | 0.424* | 0.570* | 0.449* |
| TMT – Alcohol | 0.882* | 0.724* | 0.806* | 0.707* | 0.953* | 0.539* |

All correlations were in a positive direction unless noted by (-). \* - $p < 0.05$

These data suggest that TMT exposure alone increases correlation between activity in the mPFC and amygdalar subregions, while blunting the effects of alcohol on BLA neuronal activity.

### 3.3. Experiment 3: Comparison of single and repeated TMT exposure on behavior and alcohol self-administration

The purpose of Experiment 3 was to build upon the results in Experiment 1 and compare the effects of single (sTMT) versus repeated TMT (rTMT) exposure on alcohol self-administration.

#### 3.3.1. TMT exposure does not change behavior in open field, elevated plus maze, or acoustic startle response tests

7 days after the final TMT exposure, rats began testing in the behavioral screens. *Open field.* There was no effect of TMT exposure on percent center time or total distance travelled (Table 1). One rat in the control group was identified as an outlier (distance travelled 2 standard deviations below the mean) and excluded from the experiment. *Elevated plus maze.* There was no effect of TMT exposure on percent open arm time or total distance travelled (Table 1). *Acoustic startle response.* There was no effect of TMT exposure on peak startle amplitude or startle habituation index (Table 1).

Consistent with Experiment 1, in which there was no change in behavior in the open field test 2 days after repeated TMT exposure, in this experiment there was no change in behavior in in the open field, elevated plus, or acoustic startle tests 1 week after single or repeated TMT exposure.

#### 3.3.2. Single but not repeated TMT increases alcohol self-administration

Examination of self-administration behavior with two-way RM ANOVAs across sucrose fading showed a main effect of reinforcer on alcohol lever responses (Fig 5A; F(17,544) = 61.1, p < 0.001), inactive lever responses (Table 2; F(17,544) = 1.66, p = 0.046), and locomotion (Table 2; F(17,544) = 4.91, p < 0.001) with no main effect of TMT or TMT by by reinforcer interaction. Examination of total alcohol intake (g/kg) across sucrose fading showed significantly greater alcohol intake in the sTMT group (Fig 5B; F(2,32) = 4.85, p = 0.015).

**Fig. 5.**
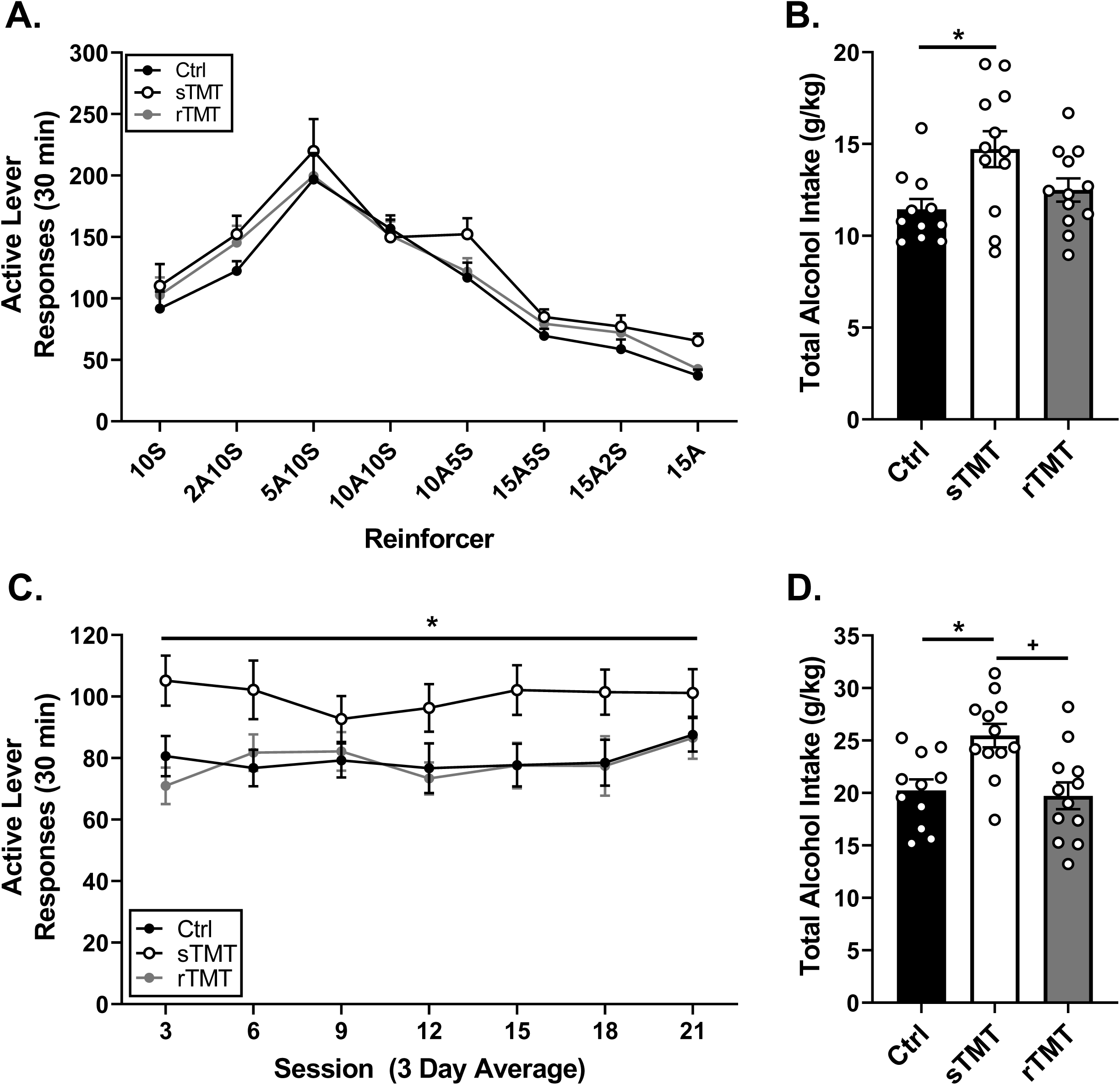
Acquisition and maintenance of alcohol self-administration following TMT exposure in Experiment 3. (A) There were no significant differences between groups in active lever responses during sucrose fading. (B) The sTMT group showed significantly higher alcohol intake than the control group during sucrose fading. (C) The sTMT group showed significantly higher active lever responses during maintenance than controls. (D) The sTMT group had significantly higher alcohol intake than both the control and rTMT groups across the maintenance phase. sTMT: single TMT exposure; rTMT: repeated TMT exposures. * - p < 0.05 versus control, + - p < 0.05 versus rTMT.

Analysis of maintenance of alcohol self-administration found a significant main effect of TMT exposure on active lever responses (Fig 5C; F(2,32) = 5.75, p = 0.007) with no significant effect of session or TMT exposure by session interaction. Again, the sTMT group consumed significantly more alcohol during the maintenance of alcohol self-administration (Fig 5D; F(2,32) = 7.70, p = 0.002). There was no effect of TMT exposure, session, or TMT exposure by session interaction on inactive lever responses (Table 3). Two-way RM-ANOVA showed a main effect of session on locomotor rate, with decreased locomotor rate across the sessions (Table 3) with no effects of TMT exposure or TMT exposure by session interaction.

These results indicate that a single TMT exposure produced lasting increase in alcohol self-administration.

### 3.4. Experiment 4: Measuring plasma corticosterone response to repeated TMT exposure

#### 3.4.1. Fecal boli production does not habituate across multiple TMT exposures

Fecal boli production during TMT exposure was measured to assess habituation of the acute physiological stress response to repeated TMT exposure (Barone et al., 1990, Monnikes et al., 1993). Two-way RM-ANOVA found a significant main effect of TMT exposure on fecal boli production (Fig 6A; F(2,32) = 20.6, p < 0.001) and a significant interaction between TMT exposure and exposure day (F(6,96) = 3.76, p = 0.002). On exposure days 1-3, the rTMT group produced significantly more fecal boli than the sTMT group (water exposure days) and Control group (p < 0.05). On the fourth exposure day both the rTMT and sTMT (TMT exposure day) groups had more fecal boli than the control group (p < 0.05). There was no habituation of fecal boli production across repeated TMT exposures.

**Fig. 6.**
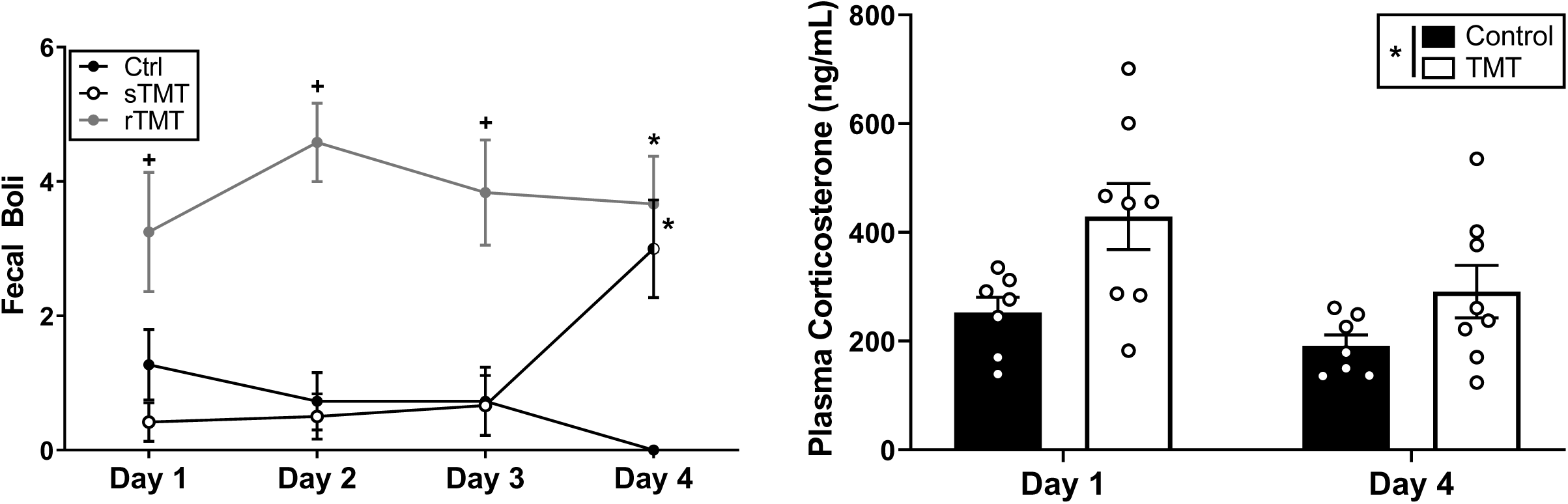
Acute physiological and endocrine stress responses to TMT across repeated exposures. (A) On days 1-3 the rTMT group produced significantly more fecal boli than the control group and sTMT group (control exposure days). On day 4 both rTMT and sTMT (TMT exposure day) groups produced significantly more fecal boli than controls. (B) Plasma corticosterone 30 minutes after exposure (at onset of dark cycle 7:00 – 9:30 pm) was significantly elevated in TMT exposed animals compared to controls on both exposure day 1 and 4. sTMT: single TMT exposure; rTMT: repeated TMT exposures. + - p < 0.05 versus control and sTMT, * - p < 0.05 versus control.

#### 3.4.2. Plasma corticosterone response does not habituate across multiple TMT exposures

In a separate group of animals, plasma corticosterone response 30 min following TMT exposure (at the onset of the dark cycle, approximately between 7:30 - 9:00 p.m.) was examined across the course of repeated TMT exposures (Fig 6B). Two-way RM-ANOVA revealed a main effect of TMT (F(1,13) = 8.74, p = 0.011) and day (F(1,13) = 5.49, p = 0.036), with no TMT by day interaction. This suggests that acute physiological and neuroendocrine responses to TMT do not habituate across repeated exposures.

## 4. Discussion

The results of this study demonstrate several important findings. First, repeated exposure to TMT, a synthetically produced predator odor, can reduce goal-tracking behavior in male Long-Evans rats. Second, rats with a history of TMT exposure show persistent increases in alcohol self-administration. Third, TMT-exposed rats are insensitive to alcohol-induced reductions in BLA c-Fos expression, and show increased synchronicity between mPFC and amygdala as indicated by c-Fos expression. Fourth, neither fecal boli nor plasma corticosterone response to TMT exposure habituate across repeated exposures. Together, these data suggest that exposure to the synthetically produced predator odor TMT is a viable model to study stress-induced lasting increases in alcohol self-administration, which may be related to changes in BLA response to alcohol.

As PTSD and addiction both involve maladaptive attribution of salience towards traumatic and drug cues, respectively, we hypothesized that stress exposure would increase the attribution of salience towards reward-related cues (i.e., increased sign-tracking behavior) as this can predict a pro-addiction phenotype (Tomie et al., 2008, Morrison et al., 2015, Fitzpatrick et al., 2019). To test this hypothesis, rats in Experiment 1 were trained on PCA, exposed to TMT and then resumed PCA. Contrary to our hypothesis, we found that sign-tracking behavior was not increased following TMT exposure, suggesting that attribution of salience towards the reward cue was not increased. In fact, we found that goal-tracking behavior was decreased following TMT exposure. It is possible that TMT exposure may still increase the salience of drug-related cues, but not sucrose-related cues, though this is unlikely as salience transfer seems well conserved across reinforcers (Morrow et al., 2011, Yager and Robinson, 2013). Another possibility is that the low sample size in this study (n = 6/condition) compounded with high variability in sign-tracking behavior (Fig 2A; Fitzpatrick et al., 2013, Fitzpatrick and Morrow, 2016) and individual differences in response to predator odor (Edwards et al., 2013, Manjoch et al., 2016, Dopfel et al., 2019) may have contributed to the lack of a group difference in sign-tracking behavior. Further, our population appeared biased towards goal-tracking behavior (Fig 2G; Fitzpatrick et al., 2013), therefore using rats from a supplier whose population demonstrates greater levels of sign-tracking behavior, or a different strain of rat may be an important consideration for future work. However, the reductions in goal-tracking behavior in the TMT-exposed rats is particularly interesting and similar reductions in reward approach behavior have been observed in rats following repeated footshock stress (Woon et al., 2019). One interpretation of this data pattern is that TMT exposure induced an anhedonia-like phenotype, a well-defined symptom of PTSD (APA, 2013, Nawijn et al., 2015, Vujanovic et al., 2017), such that a decrease in sucrose reward led to a decrease in goal-tracking behavior in the task. However, during self-administration training the TMT-exposed animals did not show reduced self-administration of a 10% sucrose solution (Fig. 4A) which makes such an explanation less likely. An alternative hypothesis is that this reduction in goal-tracking may be due to a cognitive impairment often associated with PTSD such as attention dysfunction (Esterman et al., 2013, Esterman et al., 2019). Future studies could directly test this by evaluating attentional processes in TMT-exposed animals with the rodent psychomotor vigilance test or five-choice serial reaction task.

Following PCA, animals began alcohol self-administration training (9 days after the last TMT exposure). Rats with a history of TMT exposure showed significant increases in alcohol self-administration during the sucrose fading and maintenance phases. These findings are consistent with literature using different models of predator odor stress and observing increases in alcohol drinking lasting 1-3 weeks (Edwards et al., 2013, Manjoch et al., 2016, Finn et al., 2018, Zoladz et al., 2018). However, here we show that the increase in self-administration persisted throughout 21 sessions (4 weeks) of maintenance self-administration which began approximately 5 weeks following the last TMT exposure.

While other studies use a variety of natural predator odors such as cat litter (Manjoch et al., 2016), rat bedding (Finn et al., 2018), or bobcat urine (Edwards et al., 2013) this is the first study to demonstrate that a synthetically produced predator odor (TMT) can persistently increase alcohol consumption. Furthermore, a recent study has also shown TMT can act as a stressor to induce reinstatement of alcohol-seeking behavior in mice (King and Becker, 2019). An advantage to using a synthetically produced predator odor such as TMT is in reproducibility, specifically because the intensity and composition of the predator odor can be controlled. Natural odorants such as bobcat urine can contain a number of different volatile odor compounds (Mattina et al., 1991), and their concentrations vary based on factors such as freshness of the urine, saturation of bedding, or diet/hydration of the animal (Apfelbach et al., 2015). Conversely, TMT as a single odor molecule, may not elicit the full range of behavioral and neurobiological responses, such as conditioned avoidance, that a natural odor might (McGregor et al., 2002, Blanchard et al., 2003). Future studies could explore alternative synthetically produced predator odors such as β-propylthietane, or a combination of odorants (Perez-Gomez et al., 2015).

To begin to understand the neurobiological basis of increased alcohol self-administration, Experiment 2 examined whether TMT-exposed rats showed differential expression of c-Fos, a marker of neuronal activity, in response to alcohol 1 week after the final TMT exposure. We examined subregions of the mPFC and amygdala, which are implicated in both PTSD, AUD, and activated by predator odor (Asok et al., 2013, Edwards et al., 2013, Hwa et al., 2019). Acute administration of alcohol (2 g/kg, i.g.) reduced c-Fos expression in the mPFC (PL and IL) regardless of whether rats had been exposed to TMT. Similarly, TMT exposure did not affect the alcohol-induced increase in c-Fos in the CeA. However, TMT-exposed rats treated with alcohol were insensitive to the alcohol-induced reductions in BLA c-Fos seen in control rats. The finding that TMT exposure prevented the alcohol-induced reduction in BLA c-Fos is particularly interesting as alcohol is known to blunt BLA response to fearful stimuli (Sripada et al., 2011) and thus hypo-responsiveness to the effects of alcohol may pose a potential mechanism underlying increased alcohol self-administration. Further, BLA hyperactivity is implicated in PTSD as well as in anxiety disorders and addiction (Patel et al., 2016, Sharp, 2017, Piggott et al., 2019) while alcohol self-administration in alcohol experienced rats is shown to reduce BLA activity (Vilpoux et al., 2009). Therefore, it is possible that if alcohol-induced reductions in BLA activity are disrupted in TMT-exposed animals, this lack of feedback could potentially drive increased alcohol drinking. Interestingly, the correlation between PL-BLA c-Fos in alcohol-treated TMT-exposed animals mirrors patterns evoked by context re-exposure in a different predator odor study (Edwards et al., 2013), and symptom-provocation in PTSD patients (Gilboa et al., 2004) supporting the hypothesis that maladaptive connectivity in these regions may underlie comorbidity of PTSD and AUD (Gilpin and Weiner, 2017).

Building on the findings of Experiment 1 showing a persistent escalation in alcohol self-administration in rats with a history of repeated TMT exposures, the goal of Experiment 3 was to determine whether a single TMT exposure would be sufficient to increase alcohol self-administration. Surprisingly, results from this experiment showed that a single TMT exposure, but not repeated TMT exposures, led to significantly elevated alcohol self-administration. Though animals’ conditioning history differed between experiments, in both cases the escalations in drinking persisted through 21 sessions (4 weeks) of maintenance self-administration, approximately 5 weeks after the last TMT exposure. A notable difference between the two experiments is that rats in Experiment 1 experienced PCA training prior to alcohol self-administration training. Therefore, that group had prior experience in the operant chambers with sucrose reward and the CS lever was the same as the active lever for later self-administration studies. While this confound was mitigated in the context of Experiment 1 as all rats received equal CS/US pairings, it could explain the divergent results in the repeated TMT groups in Experiments 1 and 3. Further, since a 1-day TMT exposure group was not tested in Experiment 1, we cannot conclude whether the repeated TMT exposure was necessary to show the subsequent increase in alcohol self-administration under those conditions. Interestingly, while repeated TMT did not elicit elevations in self-administration in Experiment 3, data from Experiments 3 and 4 show that neither the acute physiological (fecal boli) nor neuroendocrine (corticosterone) responses habituated across repeated TMT exposures. This is consistent with another study showing that corticosterone response of mice to repeated exposures to rat-bedding does not habituate (Finn et al., 2018), and stands in contrast to other modes of repeated stress such as restraint where corticosterone responses habituate (Schmidt et al., 2019). This data pattern highlights the unique nature of PO as a stressor evocative of an evolutionarily engrained response (Perez-Gomez et al., 2015).

In both Experiments 1 and 3 rats with a history of TMT exposure showed increased alcohol self-administration. One explanation is that the TMT exposure may have led to an enhancement in the reinforcing value of alcohol. Alternatively, the increase in self-administration could be interpreted as a reduction in the reinforcing effects of alcohol such that rats required more alcohol. Future studies could further investigate the mechanisms underlying increased self-administration by examining performance on a progressive ratio test for an alcohol reinforcer (Stafford et al., 1998). Another possible explanation is that rats were less sensitive to the interoceptive cues of alcohol that signal satiety. Indeed, prior work has shown that rats with a history of corticosterone exposure in the drinking water show blunted sensitivity to the interoceptive effects of alcohol and also show enhanced alcohol self-administration (Besheer et al., 2012, Besheer et al., 2013, Besheer et al., 2014, Jaramillo et al., 2015). One caveat to the present findings of increased alcohol self-administration is that it is not known if these effects are specific to an alcohol reward or to reward in general. As such, it will be important for future work to conduct a parallel experiment in rats trained to self-administer a non-drug reward such as sucrose or a non-caloric reward such as saccharine.

One question that remains unanswered by the existing literature is the effect that alcohol history has on the outcomes of stress exposure on alcohol drinking. Animals that showed increases in alcohol self-administration in this study were alcohol naïve, but two studies show that a history of alcohol consumption is necessary for elevations in alcohol consumption following PO or predator and social stress (Finn et al., 2018, Zoladz et al., 2018). Another study found that alcohol experience was protective against increases in alcohol consumption following SEFL (Meyer et al., 2013). The impact of alcohol history on stress-induced escalations in drinking will be an important point to address for clinical translation of these studies as alcohol use is prevalent among adults (SAMHSA, 2018) and clinical data suggests that alcohol dependence blunts cortisol stress response (Sinha et al., 2011).

It is particularly interesting that 7 days after TMT exposure, there were no differences in anxiety-like behavior, or hyperarousal relative to controls. This is in contrast to other studies using TMT that show the presence of anxiety-like behavior and hyperarousal 7 - 9 days following predator odor exposure (Brodnik et al., 2017, Schwendt et al., 2018). The lack of anxiety-like behavior or hyperarousal in this study could be due to strain differences, a floor effect, or timing of behavioral testing. Both studies showing hyperarousal and anxiety-like behavior in the acoustic startle and elevated plus maze use male Sprague-Dawley rats (Brodnik et al., 2017, Schwendt et al., 2018), so it is possible that this lack of effect might be specific to Long-Evans rats. Additionally, the absence of a difference in elevated plus maze performance may have been masked by a floor effect. For example, rats in the control group spent a low proportion of overall time in the open arms of the maze (4.25%), making it difficult to observe further reductions in open arm time. This may have been due to lighting in the testing room or testing during the light cycle, though this does not explain the lack of effect in the acoustic startle test. Furthermore, while another study reporting anxiety-like behavior in the elevated plus maze following predator odor stress shows immediate changes in alcohol consumption (Manjoch et al., 2016), the changes in self-administration in this study do not emerge until 2 weeks after TMT exposure. Therefore, it is possible that hyperarousal and anxiety-like behavior would not emerge until 2 weeks after PO exposure. Future studies could address these limitations by testing for behavioral sequelae at multiple timepoints, with a greater variety of behavioral tests, or in a different strain of rats.

Clinical studies suggest higher incidence of PTSD in females than males (Bangasser and Valentino, 2014). Therefore, a limitation of the present work is the lack of inclusion of female rats. One study that examined sex differences in alcohol consumption following predator odor exposure found that only low-drinking female rats increased alcohol consumption (Finn et al., 2018), suggesting that the mechanism by which stress impacts alcohol drinking may be different between sexes. This is supported by data from the single prolonged stress (SPS) model showing potentiation of alcohol-induced inhibition of basolateral amygdala neurons in female rats but not males following stress exposure (Ornelas and Keele, 2018). Other studies have shown sex differences in gene expression following predator odor stress, with females but not males showing upregulations in PFC and hippocampal p450scc, an enzyme responsible for synthesis of neuroactive steroids that can regulate alcohol drinking (Devaud et al., 2018), and males but not females showing changes in hippocampal CREB and ERK (Homiack et al., 2017). These observed sex differences suggest the possibility for sex differences in the findings of the present work and it will be important for future work to include female rats.

In summary, this study outlines a model by which a single exposure to a synthetically produced predator odor, TMT, can induce a persistent increase in alcohol self-administration in male Long Evans rats. Rats exposed to TMT also show insensitivity to the effects of alcohol on BLA c-Fos expression, which may suggest a dysregulation that could underlie these increases in alcohol self-administration. The acute physiological (fecal boli) and corticosterone response to repeated TMT exposure did not habituate, though repeated TMT exposure only increased alcohol self-administration in rats trained in PCA. Future studies could use this model and others to better define the effects of alcohol experience on traumatic stress reactivity, identify stress induced changes in mPFC-amygdala circuitry that may underlie escalations in drinking, and characterize sex differences in models of stress enhanced alcohol consumption. Through breadth of experimental methodologies and careful dissection of variables involved in trauma exposure, studies such as this can aid in understanding the highly variable nature of PTSD and its relationship with AUD.

## References

APA (2013) Diagnostic and statistical manual of mental disorders. 5th ed. Washington, DC: American Psychiatric Association.

Apfelbach R, Soini HA, Vasilieva NY, Novotny MV (2015) Behavioral responses of predator-naive dwarf hamsters (Phodopus campbelli) to odor cues of the European ferret fed with different prey species. Physiol Behav 146:57–66.

Asok A, Ayers LW, Awoyemi B, Schulkin J, Rosen JB (2013) Immediate early gene and neuropeptide expression following exposure to the predator odor 2,5-dihydro-2,4,5-trimethylthiazoline (TMT). Behav Brain Res 248:85–93.

Bangasser DA, Valentino RJ (2014) Sex differences in stress-related psychiatric disorders: neurobiological perspectives. Front Neuroendocrinol 35:303–319.

Barone FC, Deegan JF, Price WJ, Fowler PJ, Fondacaro JD, Ormsbee HS, 3rd (1990) Cold-restraint stress increases rat fecal pellet output and colonic transit. Am J Physiol 258:G329–337.

Besheer J, Fisher KR, Grondin JJ, Cannady R, Hodge CW (2012) The effects of repeated corticosterone exposure on the interoceptive effects of alcohol in rats. Psychopharmacology (Berl) 220:809–822.

Besheer J, Fisher KR, Jaramillo AA, Frisbee S, Cannady R (2014) Stress hormone exposure reduces mGluR5 expression in the nucleus accumbens: functional implications for interoceptive sensitivity to alcohol. Neuropsychopharmacology 39:2376–2386.

Besheer J, Fisher KR, Lindsay TG, Cannady R (2013) Transient increase in alcohol self-administration following a period of chronic exposure to corticosterone. Neuropharmacology 72:139–147.

Blaine SK, Sinha R (2017) Alcohol, stress, and glucocorticoids: From risk to dependence and relapse in alcohol use disorders. Neuropharmacology 122:136–147.

Blanchard DC, Markham C, Yang M, Hubbard D, Madarang E, Blanchard RJ (2003) Failure to produce conditioning with low-dose trimethylthiazoline or cat feces as unconditioned stimuli. Behav Neurosci 117:360–368.

Brodnik ZD, Black EM, Clark MJ, Kornsey KN, Snyder NW, Espana RA (2017) Susceptibility to traumatic stress sensitizes the dopaminergic response to cocaine and increases motivation for cocaine. Neuropharmacology 125:295–307.

Devaud LL, Alavi M, Jensen JP, Helms ML, Nipper MA, Finn DA (2018) Sexually divergent changes in select brain proteins and neurosteroid levels after a history of ethanol drinking and intermittent PTSD-like stress exposure in adult C57BL/6J mice. Alcohol.

Dopfel D, Perez PD, Verbitsky A, Bravo-Rivera H, Ma Y, Quirk GJ, Zhang N (2019) Individual variability in behavior and functional networks predicts vulnerability using an animal model of PTSD. Nat Commun 10:2372.

Edwards S, Baynes BB, Carmichael CY, Zamora-Martinez ER, Barrus M, Koob GF, Gilpin NW (2013) Traumatic stress reactivity promotes excessive alcohol drinking and alters the balance of prefrontal cortex-amygdala activity. Transl Psychiatry 3:e296.

Esterman M, DeGutis J, Mercado R, Rosenblatt A, Vasterling JJ, Milberg W, McGlinchey R (2013) Stress-related psychological symptoms are associated with increased attentional capture by visually salient distractors. J Int Neuropsychol Soc 19:835–840.

Esterman M, Fortenbaugh FC, Pierce ME, Fonda JR, DeGutis J, Milberg W, McGlinchey R (2019) Trauma-related psychiatric and behavioral conditions are uniquely associated with sustained attention dysfunction. Neuropsychology 33:711–724.

Fenster RJ, Lebois LAM, Ressler KJ, Suh J (2018) Brain circuit dysfunction in post-traumatic stress disorder: from mouse to man. Nat Rev Neurosci 19:535–551.

Finn DA, Helms ML, Nipper MA, Cohen A, Jensen JP, Devaud LL (2018) Sex differences in the synergistic effect of prior binge drinking and traumatic stress on subsequent ethanol intake and neurochemical responses in adult C57BL/6J mice. Alcohol 71:33–45.

Fitzpatrick CJ, Geary T, Creeden JF, Morrow JD (2019) Sign-tracking behavior is difficult to extinguish and resistant to multiple cognitive enhancers. Neurobiol Learn Mem 163:107045.

Fitzpatrick CJ, Gopalakrishnan S, Cogan ES, Yager LM, Meyer PJ, Lovic V, Saunders BT, Parker CC, Gonzales NM, Aryee E, Flagel SB, Palmer AA, Robinson TE, Morrow JD (2013) Variation in the form of Pavlovian conditioned approach behavior among outbred male Sprague-Dawley rats from different vendors and colonies: sign-tracking vs. goal-tracking. PLoS One 8:e75042.

Fitzpatrick CJ, Morrow JD (2016) Pavlovian Conditioned Approach Training in Rats. J Vis Exp e53580.

Gilboa A, Shalev AY, Laor L, Lester H, Louzoun Y, Chisin R, Bonne O (2004) Functional connectivity of the prefrontal cortex and the amygdala in posttraumatic stress disorder. Biol Psychiatry 55:263–272.

Gilpin NW, Weiner JL (2017) Neurobiology of comorbid post-traumatic stress disorder and alcohol-use disorder. Genes Brain Behav 16:15–43.

Homiack D, O’Cinneide E, Hajmurad S, Barrileaux B, Stanley M, Kreutz MR, Schrader LA (2017) Predator odor evokes sex-independent stress responses in male and female Wistar rats and reduces phosphorylation of cyclic-adenosine monophosphate response element binding protein in the male, but not the female hippocampus. Hippocampus 27:1016–1029.

Hwa LS, Neira S, Pina MM, Pati D, Calloway R, Kash TL (2019) Predator odor increases avoidance and glutamatergic synaptic transmission in the prelimbic cortex via corticotropin-releasing factor receptor 1 signaling. Neuropsychopharmacology 44:766–775.

Jacobsen LK, Southwick SM, Kosten TR (2001) Substance use disorders in patients with posttraumatic stress disorder: a review of the literature. Am J Psychiatry 158:1184–1190.

Jaramillo AA, Randall PA, Frisbee S, Fisher KR, Besheer J (2015) Activation of mGluR2/3 following stress hormone exposure restores sensitivity to alcohol in rats. Alcohol 49:525–532.

Jaramillo AA, Randall PA, Stewart S, Fortino B, Van Voorhies K, Besheer J (2018) Functional role for cortical-striatal circuitry in modulating alcohol self-administration. Neuropharmacology 130:42–53.

Kessler RC, Crum RM, Warner LA, Nelson CB, Schulenberg J, Anthony JC (1997) Lifetime co-occurrence of DSM-III-R alcohol abuse and dependence with other psychiatric disorders in the National Comorbidity Survey. Arch Gen Psychiatry 54:313–321.

King CE, Becker HC (2019) Oxytocin attenuates stress-induced reinstatement of alcohol seeking behavior in male and female mice. Psychopharmacology (Berl) 236:2613–2622.

Koob GF, Volkow ND (2016) Neurobiology of addiction: a neurocircuitry analysis. Lancet Psychiatry 3:760–773.

Liberzon I, Sripada CS (2008) The functional neuroanatomy of PTSD: a critical review. Prog Brain Res 167:151–169.

Makhijani VH, Van Voorhies K, Besheer J (2018) The mineralocorticoid receptor antagonist spironolactone reduces alcohol self-administration in female and male rats. Pharmacol Biochem Behav 175:10–18.

Manjoch H, Vainer E, Matar M, Ifergane G, Zohar J, Kaplan Z, Cohen H (2016) Predator-scent stress, ethanol consumption and the opioid system in an animal model of PTSD. Behav Brain Res 306:91–105.

Mattina MJ, Pignatello JJ, Swihart RK (1991) Identification of volatile components of bobcat (Lynx rufus) urine. J Chem Ecol 17:451–462.

McGregor IS, Schrama L, Ambermoon P, Dielenberg RA (2002) Not all ‘predator odours’ are equal: cat odour but not 2,4,5 trimethylthiazoline (TMT; fox odour) elicits specific defensive behaviours in rats. Behav Brain Res 129:1–16.

Meyer EM, Long V, Fanselow MS, Spigelman I (2013) Stress increases voluntary alcohol intake, but does not alter established drinking habits in a rat model of posttraumatic stress disorder. Alcohol Clin Exp Res 37:566–574.

Monnikes H, Schmidt BG, Tache Y (1993) Psychological stress-induced accelerated colonic transit in rats involves hypothalamic corticotropin-releasing factor. Gastroenterology 104:716–723.

Morrison SE, Bamkole MA, Nicola SM (2015) Sign Tracking, but Not Goal Tracking, is Resistant to Outcome Devaluation. Front Neurosci 9:468.

Morrow JD, Maren S, Robinson TE (2011) Individual variation in the propensity to attribute incentive salience to an appetitive cue predicts the propensity to attribute motivational salience to an aversive cue. Behav Brain Res 220:238–243.

Nawijn L, van Zuiden M, Frijling JL, Koch SB, Veltman DJ, Olff M (2015) Reward functioning in PTSD: a systematic review exploring the mechanisms underlying anhedonia. Neurosci Biobehav Rev 51:189–204.

Ornelas LC, Keele NB (2018) Sex Differences in the Physiological Response to Ethanol of Rat Basolateral Amygdala Neurons Following Single-Prolonged Stress. Front Cell Neurosci 12:219.

Patel R, Girard TA, Pukay-Martin N, Monson C (2016) Preferential recruitment of the basolateral amygdala during memory encoding of negative scenes in posttraumatic stress disorder. Neurobiol Learn Mem 130:170–176.

Perez-Gomez A, Bleymehl K, Stein B, Pyrski M, Birnbaumer L, Munger SD, Leinders-Zufall T, Zufall F, Chamero P (2015) Innate Predator Odor Aversion Driven by Parallel Olfactory Subsystems that Converge in the Ventromedial Hypothalamus. Curr Biol 25:1340–1346.

Piggott VM, Bosse KE, Lisieski MJ, Strader JA, Stanley JA, Conti AC, Ghoddoussi F, Perrine SA (2019) Single-Prolonged Stress Impairs Prefrontal Cortex Control of Amygdala and Striatum in Rats. Front Behav Neurosci 13:18.

Randall PA, Stewart RT, Besheer J (2017) Sex differences in alcohol self-administration and relapse-like behavior in Long-Evans rats. Pharmacol Biochem Behav 156:1–9.

Rau V, Fanselow MS (2009) Exposure to a stressor produces a long lasting enhancement of fear learning in rats. Stress 12:125–133.

SAMHSA SAaMHA (2018) 2018 National Survey on Drug Use and Health (NSDUH)

Schmidt KT, Makhijani VH, Boyt KM, Cogan ES, Pati D, Pina MM, Bravo IM, Locke JL, Jones SR, Besheer J, McElligott ZA (2019) Stress-Induced Alterations of Norepinephrine Release in the Bed Nucleus of the Stria Terminalis of Mice. ACS Chem Neurosci 10:1908–1914.

Schwendt M, Shallcross J, Hadad NA, Namba MD, Hiller H, Wu L, Krause EG, Knackstedt LA (2018) A novel rat model of comorbid PTSD and addiction reveals intersections between stress susceptibility and enhanced cocaine seeking with a role for mGlu5 receptors. Transl Psychiatry 8:209.

Seo D, Sinha R (2014) The neurobiology of alcohol craving and relapse. Handb Clin Neurol 125:355–368.

Sharp BM (2017) Basolateral amygdala and stress-induced hyperexcitability affect motivated behaviors and addiction. Transl Psychiatry 7:e1194.

Shorter D, Hsieh J, Kosten TR (2015) Pharmacologic management of comorbid post-traumatic stress disorder and addictions. Am J Addict 24:705–712.

Sinha R, Fox HC, Hong KI, Hansen J, Tuit K, Kreek MJ (2011) Effects of adrenal sensitivity, stress- and cue-induced craving, and anxiety on subsequent alcohol relapse and treatment outcomes. Arch Gen Psychiatry 68:942–952.

Sripada CS, Angstadt M, McNamara P, King AC, Phan KL (2011) Effects of alcohol on brain responses to social signals of threat in humans. Neuroimage 55:371–380.

Stafford D, LeSage MG, Glowa JR (1998) Progressive-ratio schedules of drug delivery in the analysis of drug self-administration: a review. Psychopharmacology (Berl) 139:169–184.

Tomie A, Grimes KL, Pohorecky LA (2008) Behavioral characteristics and neurobiological substrates shared by Pavlovian sign-tracking and drug abuse. Brain Res Rev 58:121–135.

Valyear MD, Villaruel FR, Chaudhri N (2017) Alcohol-seeking and relapse: A focus on incentive salience and contextual conditioning. Behav Processes 141:26–32.

VanElzakker MB, Dahlgren MK, Davis FC, Dubois S, Shin LM (2014) From Pavlov to PTSD: the extinction of conditioned fear in rodents, humans, and anxiety disorders. Neurobiol Learn Mem 113:3–18.

Vilpoux C, Warnault V, Pierrefiche O, Daoust M, Naassila M (2009) Ethanol-sensitive brain regions in rat and mouse: a cartographic review, using immediate early gene expression. Alcohol Clin Exp Res 33:945–969.

Vujanovic AA, Wardle MC, Smith LJ, Berenz EC (2017) Reward functioning in posttraumatic stress and substance use disorders. Curr Opin Psychol 14:49–55.

Woon EP, Seibert TA, Urbanczyk PJ, Ng KH, Sangha S (2019) Differential effects of prior stress on conditioned inhibition of fear and fear extinction. Behav Brain Res 381:112414.

Yager LM, Robinson TE (2013) A classically conditioned cocaine cue acquires greater control over motivated behavior in rats prone to attribute incentive salience to a food cue. Psychopharmacology (Berl) 226:217–228.

Zoladz PR, Eisenmann ED, Rose RM, Kohls BA, Johnson BL, Robinson KL, Heikkila ME, Mucher KE, Huntley MR (2018) Predator-based psychosocial stress model of PTSD differentially influences voluntary ethanol consumption depending on methodology. Alcohol 70:33–41.

